# SOLeNNoID: A Deep Learning Pipeline For Solenoid Residue Detection in Protein Structures

**DOI:** 10.1101/2024.07.22.604558

**Authors:** Georgi Nikov, Daniella Pretorius, James W. Murray

**Affiliations:** Department of Life Sciences, Imperial College London, Exhibition Road, London SW7 2AZ, United Kingdom

## Abstract

Solenoid proteins are a subset of tandem repeat proteins, which are structurally distinct from globular proteins. Solenoid proteins are defined by their modular, elongated structures, dependent on interactions between adjacent repeats. These proteins are found across all domains of life and have many important functions such as protein binding, enzymatic catalysis, ice binding and nucleic acid binding. Furthermore, engineered variants of solenoid proteins such as DARPins and designed PPR proteins have therapeutic commercial applications. In order to advance the study of natural solenoid proteins and the design of novel solenoid proteins, accurate tools for solenoid detection and annotation are required. As solenoid structures are more conserved than solenoid sequences and owing to recent developments in protein structure prediction, structure-based solenoid detection is preferred. Here we propose SOLeNNoID - a deep learning pipeline for solenoid residue prediction in protein structures. We cover all three solenoid sub-classes: alpha-, alpha/beta- and beta-solenoids. We use a CNN architecture to reason over protein distance matrices and compare our method to existing structure-based methods. Finally, we produce predictions on the entire PDB and demonstrate a 71 percent increase in solenoid-containing entries over the gold-standard RepeatsDB database using our method.

**GitHub:** https://github.com/gnik2018/SOLeNNoID

**Zenodo:** TBC

## Introduction

Tandem repeat proteins (TRPs) consist of multiple identical or highly similar structural modules and are defined primarily by intra-repeat and inter-repeat interactions between residues adjacent in the primary sequence (1) as opposed to the longer-range interactions between residues distant in the primary sequence observed in globular proteins. It is estimated that about 50 percent of proteins contain at least one tandem repeat, defined as an adjacently repeated amino acid pattern (2). Tandem repeat proteins can be divided into five main classes (3). Class I consists of short (1-to-2-residue) repeats which form crystallites, whereas class II comprises fibrous repeating structures such as collagen and alpha-helical coiled coils. Class III includes solenoid and non-solenoid ‘open’ structures, characterised by a modular elongated structure which is maintained by inter-repeat interactions. Class IV comprises modular ‘closed’ structures where the N- and C-terminal repeats of the protein interact. Class V is made up of beads-on-a-string proteins, where each repeat is a small independently folded domain connected to adjacent repeats via linker peptides.

Solenoid proteins (class III) are structurally and functionally diverse. For example, solenoid proteins can bind to nucleic acids (4), catalyse reactions (5), display antifreeze activity (6) and bind to ligands (7). In addition, solenoid proteins are attractive design targets due to their modular architecture and have commercial applications in RNA editing (pentatricopeptide repeat proteins - EditForce) and cancer therapy (DARPins - Molecular Partners), in addition to numerous binding and materials applications developed by academic groups (8). The polypeptide chain in solenoid structures follows a helical path, where individual repeats pack against each other and depend on each other for folding (1). Individual repeats (one turn around the solenoid axis) are made up of secondary structure elements ranging from alpha-helices and beta-strands to 3_10_ and polyproline II helices connected by loops.

Both the design of novel solenoid proteins and the exploration of existing ones rely on the accurate detection of repeating units, repeating regions and entire domains. Protein sequence information is much more abundant than experimental structural information. For example, the 03 Aug 2022 release of the UniProtKB/TrEMBL protein sequence database (9) contains 226,771,949 entries, while, at the time of writing, the PDB (10, 11) has only 222,624 entries, over a thousand times fewer. Sequence-based tandem repeat detection approaches range from using self-alignment matrices (e.g. TRUST (12)), k-means clustering (T-REKS (13)), Hidden Markov Model (HHRepID (14)), and discrete Fourier transform (REPETITA (15)), to neuralnetwork-based approaches (16–19). In addition, the TRAL library (20, 21) integrates many of the above tandem repeat detectors as well as post-processing, refinement, and annotation modules to evaluate and improve the detected repeats. Sequence-level tandem repeat detection is more challenging than with structures due to deviations from ideality in tandem repeat sequences such as insertions within and between repeats, and sequence divergence between individual repeats (22). Structural detection of repeats has become more applicable recently, as large databases of accurate structure predictions such as AlphaFold (23) and ESMfold (24) have become available. Examples of recent structural tandem repeat detection methods include TAPO (25), RepeatsDB-lite (26) and PRIGSA2 (22). TAPO comprises an ensemble of 7 submethods, which use features such as periodicities of atomic coordinates, conformational alphabet strings, residue contact maps and secondary structure vectors. The outputs of these sub-methods are combined using a support vector machine (SVM) model, to detect tandem repeats. RepeatsDB-lite relies on iteratively matching tandem repeats from a predefined library to the input structure to expand the predicted repeating region. PRIGSA2 is a graph-based algorithm, which analyses the contact network and secondary structure annotation of proteins to identify tandem repeat units.

While many tandem repeat prediction algorithms have been developed, there have been relatively few attempts to leverage neural networks. Convolutional neural networks (CNNs) have been applied to detect tandem repeat proteins using 3D protein structure data (DeepSymmetry (18)) and 2D distance matrix representations of protein structure (Deep-StRIP (19)). Deep-StRIP uses two CNN models - one which classifies a protein as belonging to class III, class IV, or non-tandem-repeat, and one which identifies the repeat region after classification. The predictions of the second model are then further processed using a smoothing algorithm.

Here we focus on the solenoid sub-classes (alpha, alpha/beta, beta) (27) of tandem repeat proteins and propose a structure-based supervised deep learning method, SOLeNNoID, for high-quality annotation of repeating regions. We use a 2D distance matrix representation of 3D protein structure and a semantic segmentation approach, employing a U-Net convolutional neural network architecture (28) to produce a probability that a given amino acid in the protein structure belongs to one of the solenoid classes, or is non-solenoid. We showcase our method and downstream analysis of predictions, compare our approach to state-of-the-art methods such as TAPO, RepeatsDB-Lite and PRIGSA2, and search the PDB for alpha-, alpha/beta- and beta-solenoid proteins to add further entries onto databases such as RepeatsDB (29) and Db-StRiPs (30).

## Methods

### Protein Structure Representation

The structure of a protein can be represented by a matrix of distances between Cα atoms (31). The distance matrix is orientation and translation invariant. Each row *n* of the distance matrix contains the distances from amino acid *x*_*n*_ to all other amino acids in the protein. Information about the structural environment of that amino acid is contained within the pattern of distances.

### Datasets

The training/validation dataset consisted of the dataset produced by Marsella et al (15) and additional beta-solenoid data. Additional beta-solenoid proteins were selected to better represent the diversity of cross-sectional shapes described by Kajava and Steven (32). The dataset comprised a set of protein PDB files and label files. The label files were produced using the annotations in the original dataset, and by annotating structures in PyMOL (33) for the additional beta-solenoid data. Each label file contained a label for each residue in the corresponding PDB file. We assigned an integer value label for each residue, corresponding to either of the four possible classes – non-solenoid (0), beta-solenoid (1), alpha/beta-solenoid (2), or alpha-solenoid (3). We assigned missing data to class 4 to distinguish from non-solenoid data. In total, the dataset consisted of 246 non-solenoid, 52 beta-solenoid, 17 alpha/beta-solenoid and 33 alpha-solenoid proteins. The training and validation datasets were derived by a 80:20 (test:validation) random split of the PDB and corresponding label files.

The test dataset consisted of 49 non-solenoid, 15 beta-solenoid, 7 alpha/beta-solenoid and 11 alpha-solenoid proteins. The non-solenoid proteins were collected from the PDBe (34)while the solenoid proteins were collected from manually reviewed solenoid entries from RepeatsDB. All structures are non-redundant with respect to the training and validation structures to a TM-align TM-score threshold of below 0.6 (35). If an alignment between two proteins produces a TM-score of above 0.5, it can be assumed that the proteins generally have the same fold in SCOP (36) or CATH (37). Thus, as the neural network will be used to identify and annotate proteins broadly in the same folds as the structures in the training dataset, the less stringent threshold of TM-score less than 0.6 was used. The RepeatsDB solenoid labels were amended by excluding loops from repeat definitions and adding parts of partial or imperfect repeats. Table S1 shows a number of statistics relating to these datasets. The datasets contain few solenoid structures but this limited data is sufficient to train a classification network. All datasets are imbalanced: there are more non-solenoid structures and residues compared to either of the solenoid classes, and there are fewer alpha/beta-solenoid structures and residues compared to the alpha- and beta-solenoid classes. For example, in the training dataset, the alpha-solenoid to non-solenoid class imbalance is 1 to 9.6, and the alpha-solenoid to alpha/beta-solenoid class imbalance is 2.2 to 1.

### Data Processing

A Cα distance matrix was calculated from each structure file using the BioPython (38) and NumPy (39) libraries in Python3 (40). At points where protein chains were discontinuous, both the labels and the distance matrix entries were replaced with padding values to maintain the register of the amino acid indices. Each distance matrix was pre- and post-padded in both x and y axes to a total length divisible by 128. Each distance matrix was then normalised by the largest distance observed in that matrix.

Per-residue class labels were converted to a 2D label matrix (or ‘segmentation mask’). Each value in the label matrix corresponds to a Cα-Cα distance, which for two residues of the same solenoid type corresponds to the solenoid type (1, 2 or 3), and is 0 otherwise. The label matrix gives labelled inter-solenoid distances. For example, consider a 70-residue protein, where residues 1-50 are beta-solenoid, and residues 51-70 are non-solenoid. The label matrix at indices from (1,1) to (50,50) would have a class value of 1, whereas the label matrix at all other indices would have a class value of 0. The label matrix was then padded identically to the distance matrix. Missing data and padding were assigned a separate class (4) to distinguish from the non-solenoid class.

Random cropping was used as a data augmentation technique to increase the amount of training data and improve model generalisation. Training distance matrix data consisted of 128×128 distance matrix slices centred on the main diagonal at a random offset. The randomly cropped distance matrix slices had corresponding slices of the label matrix as labels. However, the label matrices were of a size 64×64 and only contained label data for the central 64×64 slice of the corresponding distance matrix. The reasoning here is that the model could learn to better classify the pixels in a 64×64 slice of the distance matrix by ‘seeing’ a larger 128×128 context. Validation set distance matrices were split into overlapping 128×128 slices with an overlap of 64×64 and without random cropping. Validation set label matrices again corresponded to the central 64×64 slice of the distance matrix. Finally, the label matrices for both training and validation sets were onehot encoded.

### Model Architecture and Training

The model code was written in Python using the Tensorflow (41), Keras (42), Scikit-learn (43) and NumPy (39) libraries. The model has a modified U-Net architecture (28) with a reduced number of filters per layer (Fig. S1). Gaussian noise was applied to the input distance matrix with a standard deviation of 0.0001. The noise was added as a regularisation measure to reduce model overfitting and resulted in modest gains in performance. The convolutional layers within each block had the same number of filters of size 3×3 and stride of 1. All convolutional layers had an exponential linear unit (ELU) activation function (44). The number of filters doubled with each subsequent convolutional block in the encoder arm of the U-Net (Fig. S1, orange) and decreased by a factor of two in the decoder arm of the U-Net (Fig. S1, green). The fraction of activations zeroed by the dropout layer is shown for each convolutional block. Max pooling and transposed convolution with a filter size of 2×2 and a stride of 2 were used. Finally, a label matrix was output by a final convolutional layer with a softmax activation function highlighting regions in the distance matrix which belong to the different classes. The final upsampling block of the U-Net was removed to yield a 64×64 label matrix prediction for each 128×128 input distance matrix slice. Early stopping and model checkpointing callbacks were used with both callbacks tracking loss over the validation dataset. Training was performed for 100 epochs (approx. 1 day) with the categorical cross-entropy loss function and a batch size of 64 using the Adam (45) optimiser with a learning rate of 0.01 on an NVIDIA P1000 GPU. The model weights leading to the lowest loss value on the validation set were saved.

### Inference and Further Analysis

For inference, an individual coordinate file and chain ID are specified. The structure (in mmCIF format) is loaded and Cα coordinates and residue indices are used to produce the Cα distance matrix. At inference, the distance matrix is processed analogously to the distance matrices in the validation set. The predicted onehot label matrices are converted to integer class labels. The class of a residue is the label matrix value at indices (x,y) where x=y=residue index.The non-solenoid and missing data classes are merged.

Predictions are mapped onto the structure and visualised using the nglview library (46), while the sequence mapping is visualised using the Bokeh library (Bokeh Development Team, 2022) in a Jupyter notebook (47). User-defined repeat splitting was achieved by recording the indices of clicked equivalent positions in the nglview viewer. The recorded indices were then used to fill in repeats from the predicted solenoid residues. These repeats were then output in PDB format. The output repeats were processed using mTM-align (48), producing a PDB file with superposed repeats, TM-score and RMSD self-similarity matrices of the repeats, and a fasta file of the aligned repeat sequences. The fasta file was used to produce a logo plot of the repeat sequence alignment with the logomaker library (49).

### Benchmark Method Predictions and Processing

For all comparison methods, predictions were collected using the respective servers (TAPO: https://bioinfo.crbm.cnrs.fr/tools/tapo/index_tapo.php; RepeatsDB-Lite: http://old.protein.bio.unipd.it/repeatsdb-lite/; PRIGSA2: https://bioinf.iiit.ac.in/PRIGSA2/) and adapted to the labelling scheme for this method. For TAPO, the top prediction was used. In contrast to SOLeNNoID, the other methods used detect individual repeats instead of individual amino acids. RepeatsDB-Lite and PRIGSA2 provide class labels, whereas TAPO does not. RepeatsDB-Lite and PRIGSA2 repeat predictions were treated as non-solenoid, if predicted to belong to an incorrect class outside the scope of SOLeNNoID (for example, if the ground truth class is alpha-solenoid but instead TIM-barrel is predicted). Precision, Recall, F1-score and multi-class Matthews correlation coefficient (MCC) were used to compare the performance of the solenoid detection algorithms. The scikit-learn classification_report was used to produce Precision, Recall and F1-score metrics and matthews_corrcoef to produce the Matthews correlation coefficient measure. Confusion matrices were produced using scikit-learn confusion_matrix and ConfusionMatrixDisplay.

### PDB Solenoid Prediction Processing

mmCIF format files from the PDB as of 30 June 2021 were used. The data collected for each structure included the 4-letter PDB ID, chain ID, the number of residues for the structure and the number, percentage, and residue indices of solenoid predictions for all classes. PDB IDs from the training/validation and test datasets were removed from the final list of processed structures to limit the analysis to new protein chains. A protein chain was considered a member of a given solenoid class, if 50 percent or more of its residues were predicted to belong to this solenoid class. SOLeNNoID predictions for the whole PDB were produced locally over approximately 3.5 days on an M1 MacBook Air with 16 GB RAM.

### RepeatsDB and DbStRiPs Solennoid Entry Selection

Data for RepeatsDB solenoid proteins were downloaded from https://www.repeatsdb.org/classification/3. Subsequently, only entries with a ‘Classification’ tab beginning with ‘3.1’, ‘3.2’, or ‘3.3’ were selected, corresponding to beta-, alpha/beta- and alpha-solenoid entries, respectively. The version of RepeatsDB accessed in this work used data from the PDB as of 14 August 2022. Data for DbStRiPs were downloaded from https://bioinf.iiit.ac.in/dbstrips/data.php, selecting Type ‘Structural class’ and Sub-type ‘Alpha Solenoids’, ‘Beta Solenoids’ and ‘Alpha-Beta Solenoids’. The version of DbStRiPs accessed in this work used data from the PDB as of 03 June 2019.

## Results

### SOLeNNoID

The SOLeNNoID analysis pipeline takes as an input a PDB structure and a chain id. A Cα distance matrix is computed from the coordinates and a label matrix is predicted using the distance patterns in the distance matrix. Finally, per-residue class labels are produced by taking the solenoid class labels from the main diagonal of the label matrix. The class labels are mapped onto the input structure and sequence for visualisation. The algorithm can process arbitrarily large structures and has a quadratic time complexity (*O*(*n*^2^)) (see Fig. S2). In a test on the 6,040 AlphaFoldDB v1 predicted protein structures for *S. cerevisiae*, predictions for structures around 2500 amino acids took less than a minute, whereas predictions for structures up to around 1200 amino acids took less than 10 seconds.

In addition to the user-friendly visualisation of solenoid predictions, the SOLeNNoID Jupyter notebook also allows interactive repeat definition. Users can quickly select equivalent residues in adjacent repeats in the nglview structure viewer, (Fig. 2 A left, green sticks). These residue selections allow the solenoid region predicted by SOLeNNoID (Fig. 2 A left, magenta ribbon) to be divided into individual repeats. The resulting repeats (Fig. 2 A right) are then output as individual PDB files and processed using mTM-align and logomaker to produce informative visualisations of repeat similarity on the structure (structural similarity matrices with RMSD (Fig. 2 B left) or TM-score (Fig. 2 B right) as the criterion and sequence (logo plot) (Fig. 2 C right) levels. For example, for PDB ID 3V4E, chain A the repeat similarity matrices indicate that repeats 1-4 form a group with high similarity between members. The sequence alignment suggests that the GWG sequence is an insertion found in only one of the repeats. Consensus residues are apparent at some of the remaining positions such as Asn at position 1 and Ile at position 20. The RepeatsDB database provides a similar output for each structure.

**Fig. 1.**
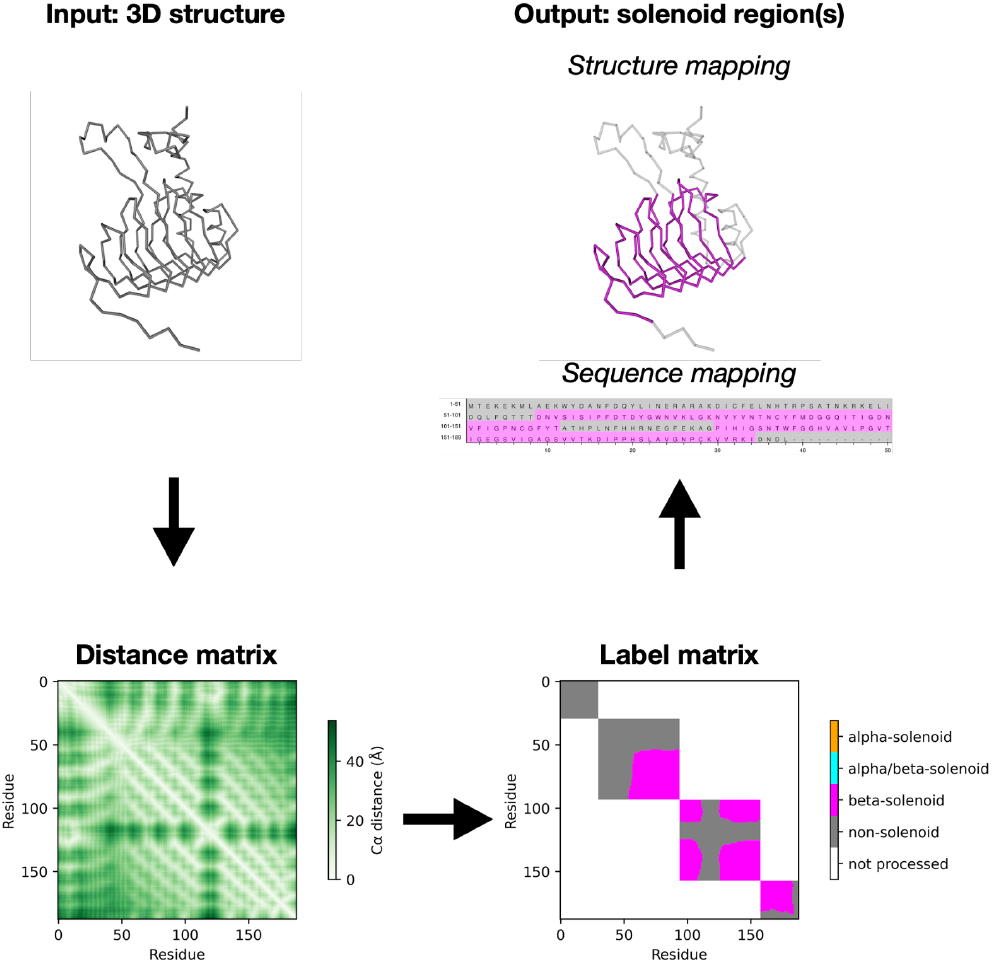
Illustration of the SOLeNNoID analysis pipeline using PDB ID 3V4E. A distance matrix is calculated from the input protein structure. A label matrix is predicted from the distance matrix using a trained U-Net neural network. Finally, the label matrix classification is converted to a per-residue classification, which is mapped onto the structure and sequence of the input protein.

**Fig. 2.**
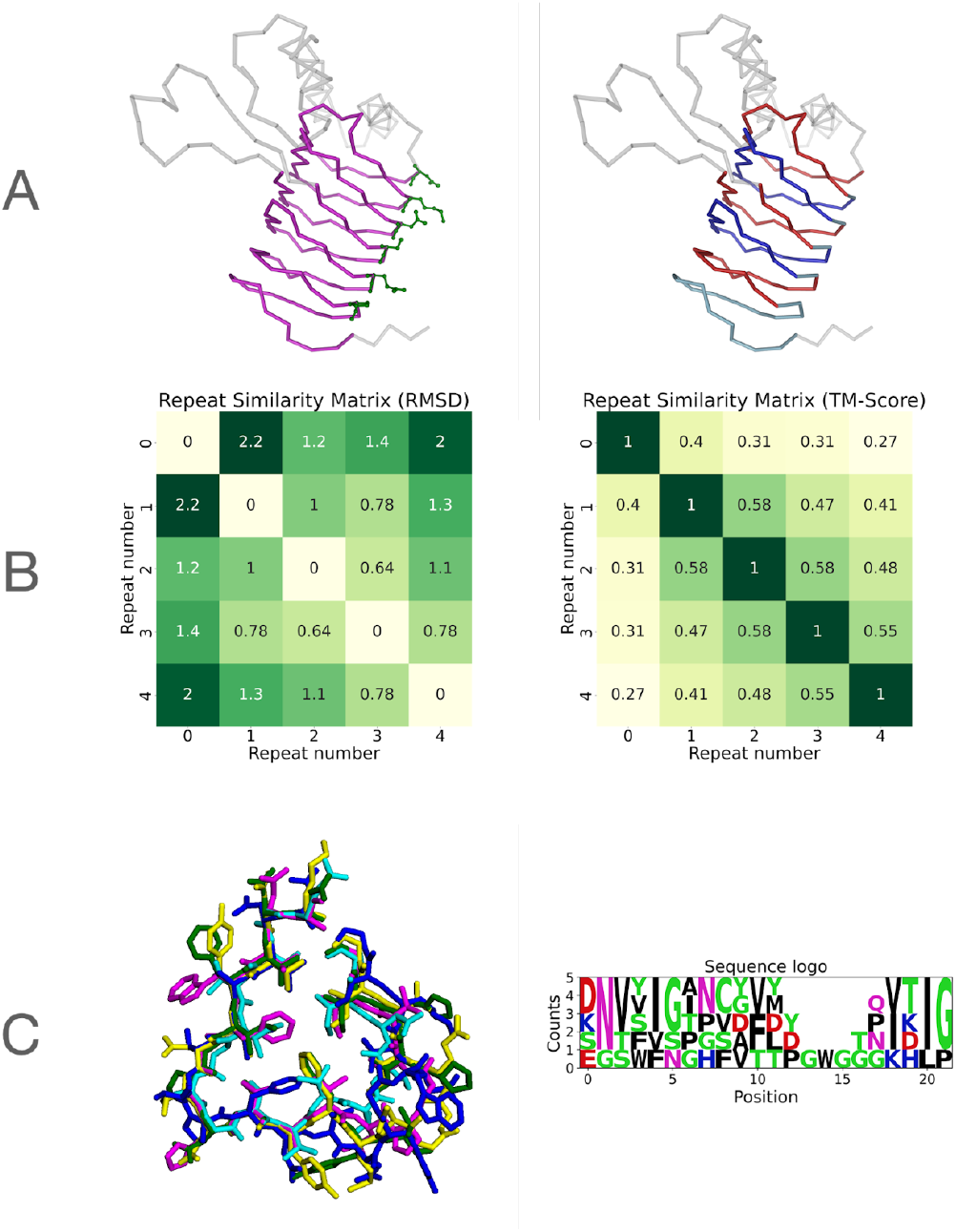
Repeat splitting and analysis output for PDB ID 3V4E chain A. A) Left: SOLeNNoID beta-solenoid predictions in magenta backbone representation and user-selected equivalent residues in each turn in green ball and stick representation. Right: Automatically defined repeats based on the user input of equivalent residues in the SOLeNNoID predictions. Alternating repeats are shown in blue and red for ease of view. B) mTM-align output repeat similarity matrices using RMSD and TM-score. C) Left: Superposition of all repeats defined from SOLeNNoID predictions and user input in line representation. Each repeat is highlighted in a different colour. Right: Sequence logo (generated using the logomaker library) for the repeats using the fasta file produced by mTM-align. Counts in the Y-axis correspond to the number of repeats of 3V4E chain A in which a residue is observed at a given position. For example, at position 0, Asp is observed in 2 repeats, while Lys, Ser and Glu are observed in 1 repeat each.

### Benchmarking against Other Tandem Repeat Detection Methods

SOLeNNoID was benchmarked against TAPO, PRIGSA2, and RepeatsDB-Lite on the solenoid structures in the test set. The four metrics used to describe performance on this dataset were precision, recall, F1-score (harmonic mean of precision and recall) and multi-class Matthews correlation coefficient (MCC). SOLeNNoID outputs solenoid residue predictions, rather than repeats. Therefore, repeat predictions of other methods were converted into a per-residue scheme. Unlike RDB-Lite and PRIGSA2, TAPO does not provide class labels. Therefore, there are two ways of comparing performance: one where it is assumed that TAPO would provide correct class labels and one where solenoid classification is treated as a binary classification of residues into a solenoid and a non-solenoid class.

If TAPO is assumed to provide correct class labels, SOLeNNoID had the best precision on the non-solenoid and beta-solenoid classes, the best recall on the alpha-solenoid class and the best F1-score on the non-solenoid and beta-solenoid classes (see 1). The PRIGSA2 method showed the highest precision on alpha/beta- and alpha-solenoid classes but had lower recall than TAPO and SOLeNNoID. Overall, SOLeNNoID outperformed PRIGSA2 and RDB-lite on F1-score and MCC and performed competitively with TAPO.

If all methods are instead used as binary solenoid/non-solenoid classifiers, SOLeNNoID outperformed TAPO in all metrics apart from recall on the solenoid class, where the two methods were equal (see Table S2). PRIGSA2 had the highest precision but very low recall on the solenoid class, ultimately resulting in a low F1-score. Therefore, while most solenoid predictions made by PRIGSA2 were true solenoid residues, many true solenoid residues were not detected. In comparison, TAPO and SOLeNNoID had a more balanced performance.

The full confusion matrices for all methods are shown in Fig. S3. Only SOLeNNoID and TAPO had large values across the entire diagonal (corresponding to residues with agreement between predicted and true label). PRIGSA2 and RDB-Lite incorrectly predicted many alpha- and beta-solenoid residues as non-solenoid. PRIGSA2 showed little to no incorrect predictions between solenoid classes. This matches the high precision but poor recall for PRIGSA2 in Table 1. RepeatsDB-Lite predicted a large number of alpha-solenoid residues as non-solenoid, which is due to some alpha-solenoid residues not being predicted as tandem repeat at all and some being predicted as a non-solenoid repeat type such as TIM-barrel or alpha-barrel. In addition, RepeatsDB-Lite incorrectly predicted many alpha/beta-solenoid residues as beta-solenoid. Both tandem repeat misclassifications by RepeatsDB-Lite are due to an incorrect classification of the master repeat unit used to build up the solenoid region. TAPO and SOLeNNoID had fewer incorrect predictions of solenoid residues as non-solenoid compared to PRIGSA2 and RepeatsDB-Lite. The major type of incorrect predictions for TAPO were non-solenoid residues predicted as beta-solenoid. SOLeNNoID instead incorrectly predicted non-solenoid residues as alpha-solenoid. This is possibly due to similarities between the distance matrices of non-solenoid alpha-helical proteins and alpha-solenoid proteins.

**Table 1.**
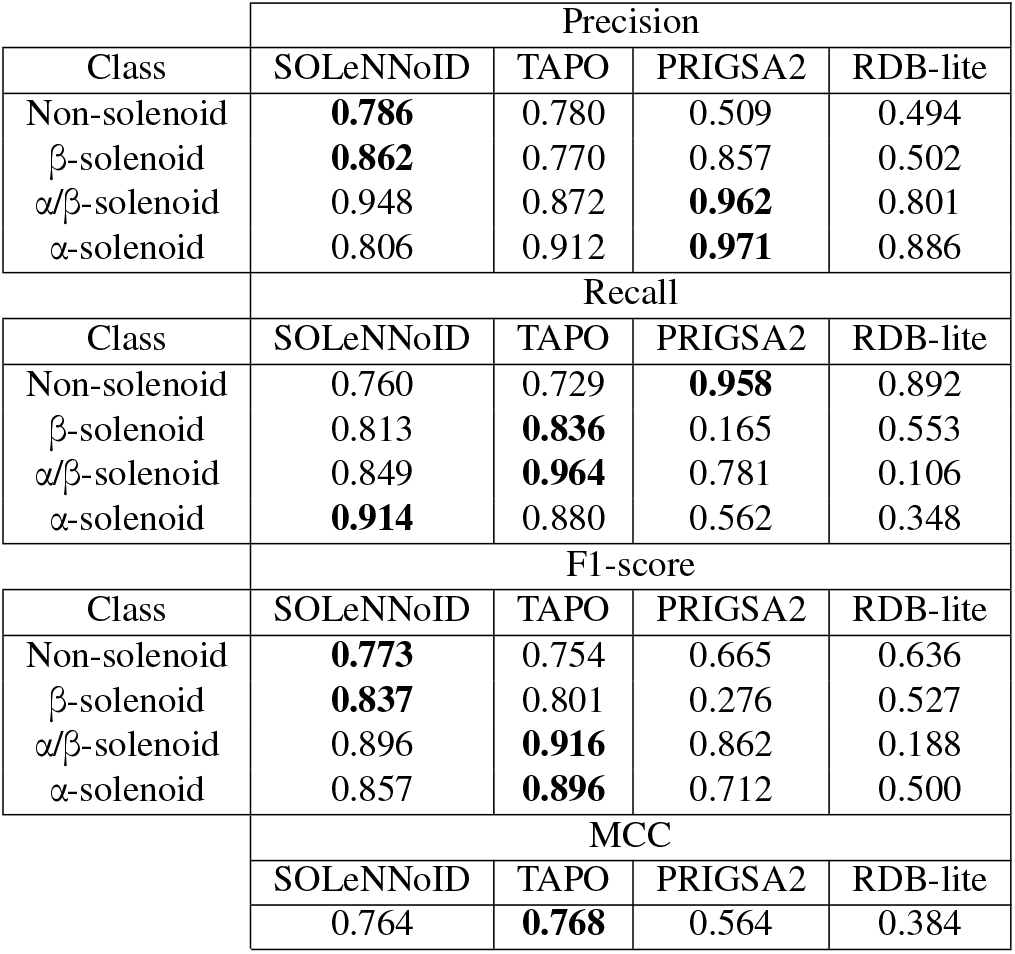
Comparison of method performance on the solenoid test set as a multi-class classification problem.

SOLeNNoID also performed well on the full test set, which includes non-solenoid structures (Table S3). On introduction of non-solenoid residues, precision and recall for the non-solenoid class increased, whereas precision for the alpha-solenoid class decreased. Precision, recall and F1-score values for solenoid classes were nearly identical to the values on the solenoid test set only apart from precision on alpha-solenoid residues, which was now lower on introduction of non-solenoid structures. This can also be seen in the bottom right corner of the confusion matrix in Fig. S4. Comparison with the confusion matrix on the solenoid test set indicates that the model incorrectly classifies an additional 256 non-solenoid residues as alpha-solenoid.

### SOLeNNoID PDB Predictions

599,443 chains (175,084 unique PDB IDs) from the PDB were processed using SOLeNNoID. Exploration of the list of alpha-solenoid predictions revealed 6 PDB IDs with over 100 chains each (3J3Q, 3J3Y, 6X63, 6QVK, 6QZ0, 6QYD). Manual inspection identified that these structures are not solenoid-containing but instead contain elements which resemble alpha-solenoid repeats. These structures were removed, leading to the final predictions outlined in Table S4. The total number of unique chains identified as containing solenoid regions by SOLeNNoID was 9,326, or about 1.56 percent of all processed PDB entries. Considering only unique PDB IDs, 4,148 PDB IDs were identified as majority solenoid, representing 2.37 percent of processed PDB IDs. This result agrees with the work of Chakrabarty and Parekh (30), who also found that approximately 2 percent of PDB entries contain solenoid regions. The total number of unique PDB IDs is lower than the sum of the numbers of unique PDB IDs of individual solenoid classes, as a single PDB ID can contain multiple solenoid chains of different classes. Therefore, using unique PDB IDs underestimates the number of solenoid structures. The mean number of solenoid residues for alpha-solenoid proteins was the highest at 348, followed by alpha/beta-solenoid proteins at 295, and beta-solenoids at 198 residues. The maximum number of solenoid residues detected in a single chain was highest for alpha-solenoid proteins (3,340: PDB ID 6TAX chain A), followed by alpha-beta-solenoid proteins (732: PDB ID 6S6Q chain B), and finally beta-solenoid proteins (631: PDB ID 6Z7P chain A). SOLeNNoID predictions also suggested that alpha-solenoid entries were by far the most abundant in the PDB. For example, there were 7.5 times more predicted alpha-solenoid chains than alpha/beta-solenoid chains.

### Comparison of PDB Predictions with Other Tandem Repeat Databases

Solenoid entries in the RepeatsDB and DbStRiPs databases had significant overlap with SOLeNNoID predictions on the PDB (see Fig. 3). There were 1,139 PDB IDs in common between RepeatsDB, DbStRiPs, and SOLeNNoID predictions, and a further 1,005 PDB IDs, which were shared between either RepeatsDB and SOLeNNoID predictions, or DbStRiPs and SOLeNNoID predictions. Therefore, SOLeNNoID predictions were supported by agreement with databases populated by distinct tandem repeat detection methods. However, SOLeNNoID also identified many PDB entries not covered by these databases. Due to detecting multiple chains from protein complexes, the number of solenoid-containing chains predicted by SOLeNNoID was about 2.6 times larger than the number of solenoid-containing PDB IDs. SOLeNNoID predictions had the second largest number of total solenoid chains (9,326) and method/database-specific solenoid chains (5,321).

**Fig. 3.**
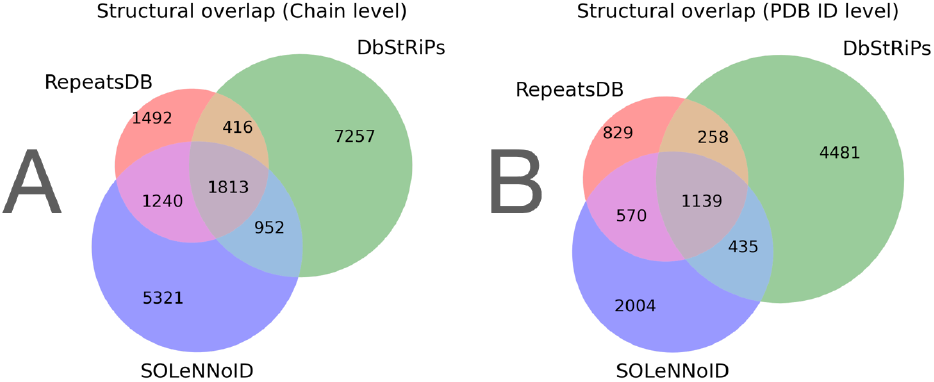
Venn diagrams showing overlap between the structures detected as solenoid by SOLeNNoID and solenoid structures in the RepeatsDB and DbStRiPs databases. Red - solenoid structures found only in the RepeatsDB database, green - solenoid structures found only in the DbStRiPs database, blue - solenoid structures found only using SOLeNNoID. Grey - solenoid structures found in all databases, pink - solenoid structures found both in RepeatsDB and by SOLeNNoID, cyan - solenoid structures found both in DbStRiPs and by SOLeNNoID, yellow - solenoid structures found both in RepeatsDB and DbStRiPs. A) Overlap between databases and SOLeNNoID predictions when considering unique individual chains. B) Overlap between databases and SOLeNNoID predictions when considering unique PDB IDs.

Table 2 shows the number of putative solenoid chains and PDB IDs found by SOLeNNoID with PDB IDs distinct from the entries in RepeatsDB and DbStRiPs. A total of 4,638 new chains and 2,004 unique PDB IDs were found with most belonging to the alpha-solenoid class. This means that 683 additional chains were discovered by SOLeNNoID for PDB IDs represented in either/both of the comparison databases. The discovery of additional solenoid structures by SOLeNNoID is due to two factors: 1) the ability of SOLeNNoID to detect examples which are not accessible to other methods, such as RepeatsDB-Lite and PRIGSA2, owing to, for example, the inability of these methods to process very large structures and 2) using a more recent version of the PDB with a larger number of chains and IDs. While the latter point applies when comparing SOLeNNoID predictions with Db-StRiPs, it does not apply to RepeatsDB, as the most recent version of the database (v3.2) is based on a version of the PDB from 2022, whereas SOLeNNoID was used to process a version of the PDB from 2021. The scale of our contribution to solenoid protein annotation is therefore demonstrated as a 71.6 percent increase over the “gold standard” RepeatsDB database. These results further highlight the need for multiple approaches to solenoid detection. Different methods show significant overlap but also complementarity and, when used together, enable a better coverage of solenoid protein structures and regions.

**Table 2.**
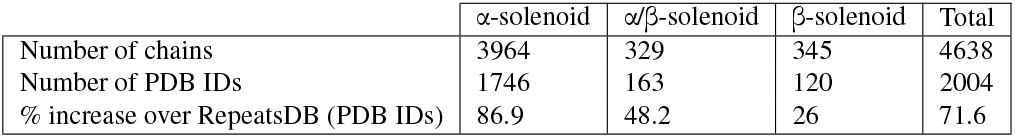
Number of structures detected by SOLeNNoID and not present in other databases and percentage increase over entries in RepeatsDB.

## Discussion

SOLeNNoID uses a CNN trained on alpha carbon distance matrices to rapidly process arbitrarily large structures and provide user-friendly output of solenoid region predictions mapped to both structure and sequence. We also present downstream interactive repeat analysis to extract structural and sequence similarities between repeats.

SOLeNNoID differs from other solenoid detection methods in that it highlights individual solenoid residues rather than repeats. Test set benchmark data shows that SOLeNNoID is competitive with TAPO and outperforms RepeatsDB-Lite and PRIGSA2. Furthermore, SOLeNNoID has balanced precision and recall. Therefore, it can be used to both annotate solenoid regions and detect new solenoid proteins, which may be overlooked if either RepeatsDB-Lite or PRIGSA2 are used. RepeatsDB-Lite depends heavily on the quality of the repeating unit library it uses to structurally align against a query protein, as well as the ‘Master’ unit, which defines the class of the repeating region. Some incorrect repeat assignments by RepeatsDB-Lite are due to incorrect ‘Master’ unit assignments - such as alpha-solenoid proteins detected as alpha-barrels, and alpha/beta-solenoid proteins detected as beta-solenoids. TAPO, in contrast, has a balanced performance as it uses several structural features in a SVM model. However, TAPO does not provide class labels. A limitation of SOLeNNoID is that it is dependent on the quality of the training dataset and solenoid proteins far outside the structural space covered by this dataset may be incorrectly predicted or omitted. All methods show certain biases on the test dataset presented here, whether predicting solenoid residues as non-solenoid, or vice versa, or confusing solenoid classes. These biases highlight the need for a consensus approach to solenoid region prediction by using the outputs of several different repeat detection methods.

Using SOLeNNoID, we have detected hundreds to thousands of new proteins with the different solenoid classes from the largest database of experimentally determined structures, the PDB. In our analysis, 50 percent of residues in a chain must be detected as solenoid - this is a stringent threshold used to remove false positive solenoid protein hits. However, this will also have the effect of under-estimating true solenoid protein numbers as solenoids with partial coverage are omitted. The PDB search for solenoid proteins we conducted revealed substantial overlap with existing databases of solenoid proteins but also complementarity of the different methods. We present over 5000 new protein chains and over 2000 new PDB IDs not covered by existing databases - a 71 percent increase over the current version of RepeatsDB. Our findings suggest that the best way to address the challenge of finding new solenoid proteins is to combine multiple approaches. With the rise of predicted structural databases with hundreds of millions of structures such as the AlphaFold and ESMFold databases, efficient analysis of structures is a key bioinformatic task. Our solenoid detection system is a contribution to analysing this flood of structural information.

## ACKNOWLEDGEMENTS

We acknowledge computational resources and support provided by the Imperial College Research Computing Service (http://doi.org/10.14469/hpc/2232). Georgi Nikov was supported by a EPSRC DTP training grant (EP/R513052/1).

## Supplementary Methods

**Table S1.**
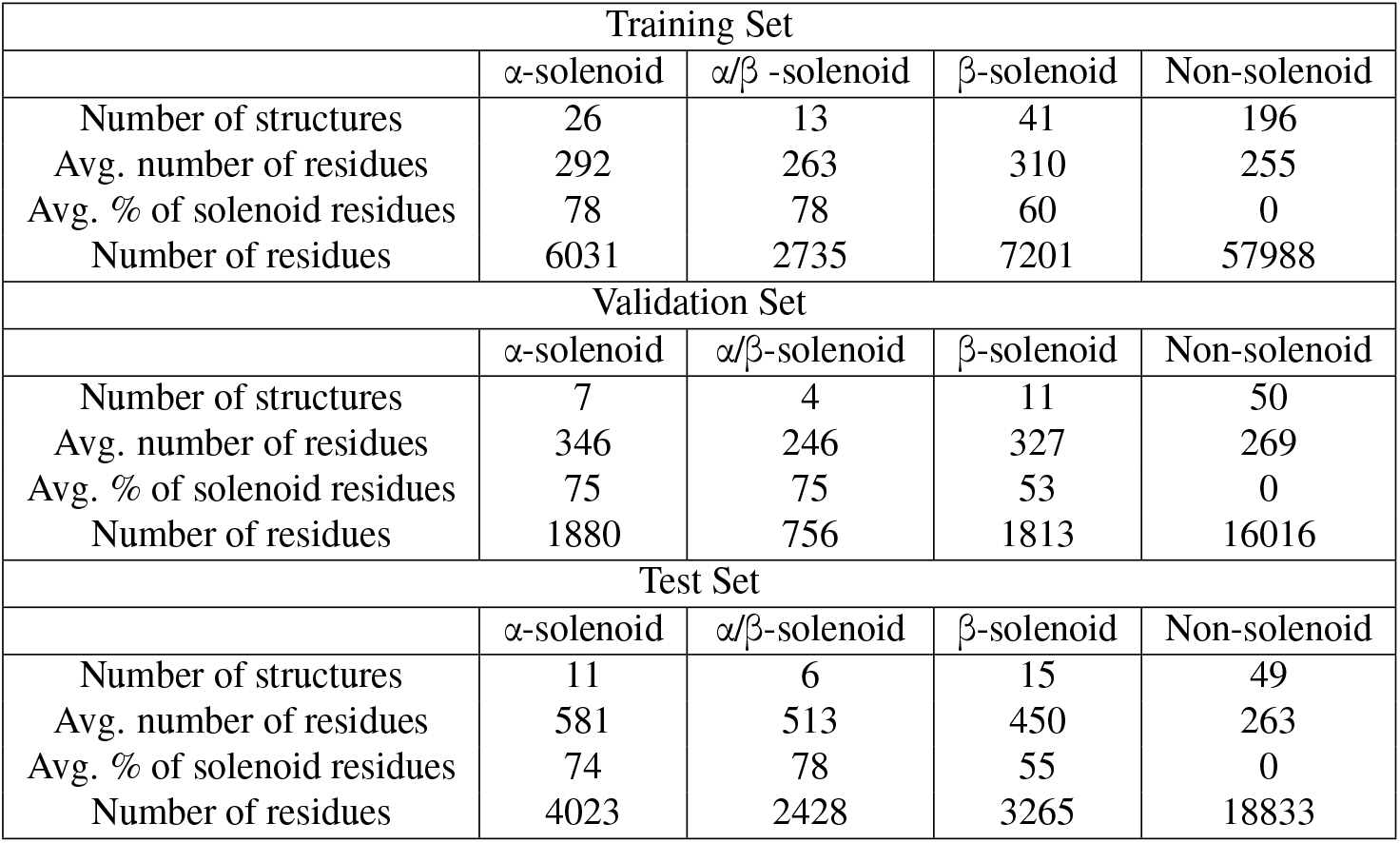
Training, validation and test set statistics.

| Training Set |  |  |  |  |
| --- | --- | --- | --- | --- |
| | α-solenoid | $\alpha/\beta$-solenoid | β-solenoid | Non-solenoid |
| Number of structures | 26 | 13 | 41 | 196 |
| Avg. number of residues | 292 | 263 | 310 | 255 |
| Avg. % of solenoid residues | 78 | 78 | 60 | 0 |
| Number of residues | 6031 | 2735 | 7201 | 57988 |
| Validation Set |  |  |  |  |
| | α-solenoid | $\alpha/\beta$-solenoid | β-solenoid | Non-solenoid |
| Number of structures | 7 | 4 | 11 | 50 |
| Avg. number of residues | 346 | 246 | 327 | 269 |
| Avg. % of solenoid residues | 75 | 75 | 53 | 0 |
| Number of residues | 1880 | 756 | 1813 | 16016 |
| Test Set |  |  |  |  |
| | α-solenoid | $\alpha/\beta$-solenoid | β-solenoid | Non-solenoid |
| Number of structures | 11 | 6 | 15 | 49 |
| Avg. number of residues | 581 | 513 | 450 | 263 |
| Avg. % of solenoid residues | 74 | 78 | 55 | 0 |
| Number of residues | 4023 | 2428 | 3265 | 18833 |

**Fig. S1.**
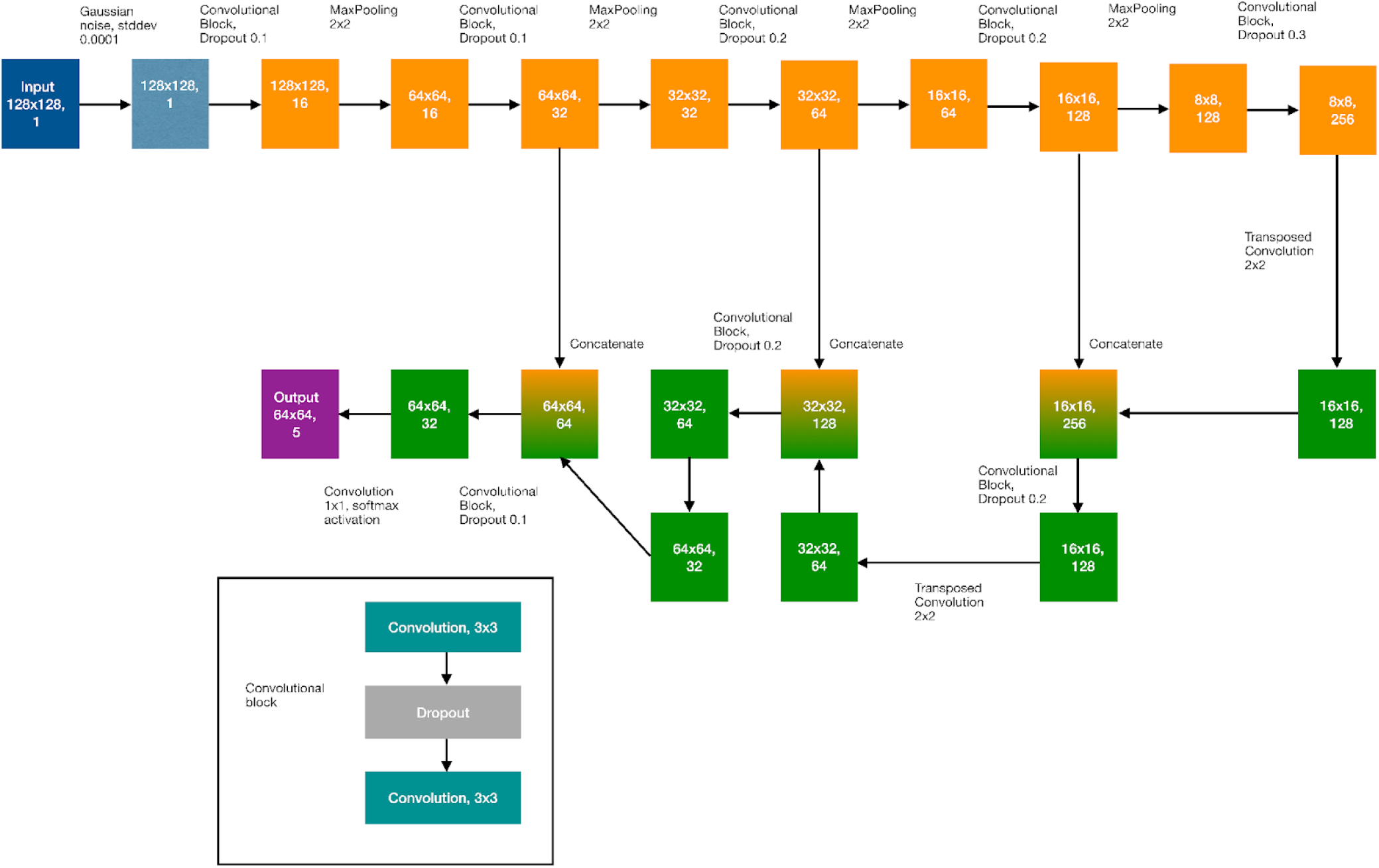
Illustration of the U-Net architecture used in this work. Each convolutional block consists of the three layers shown in the box. Each square denotes the dimensions of the input, intermediate tensor or output.

### Prediction Time Test

The prediction time test was conducted using the 6,040 AlphaFold2-predicted S. cerevisiae structures (v1 database) in the EBI AlphaFold database. The script was executed on the Imperial College HPC with 16 CPU cores, 96 GB RAM and 4 RTX6000 GPUs. The prediction time was defined from before loading a structure to after obtaining solenoid predictions for the structure.

### Data Processing and Visualisation

Data analysis was carried out using the Pandas library (50) in Python3 (40). Data visualisation was carried out using the Seaborn (51), matplotlib (52), matplotlib-venn (https://github.com/konstantint/matplotlibvenn) and logomaker (49) libraries. Protein structure visualisation was carried out using the nglview library (46) in Python3.

## Supplementary Results

### Prediction Time Test

To evaluate the time taken by SOLeNNoID to process a structure and make predictions, a test was conducted on the 6,040 v1 AlphaFoldDB predicted protein structures for S. cerevisiae. The result in Fig. S3 below shows that prediction time had a second order polynomial relationship with respect to the length of a protein. However, predictions for structures around 2500 amino acids took less than a minute under the conditions used, whereas predictions for structures up to around 1200 amino acids took less than 10 seconds. This demonstrates that SOLeNNoID can rapidly produce predictions even for large structures. In comparison, the TAPO, PRIGSA2 and RepeatsDB-Lite servers can take on the order of minutes to produce a prediction for a structure over 1000 amino acids.

**Fig. S2.**
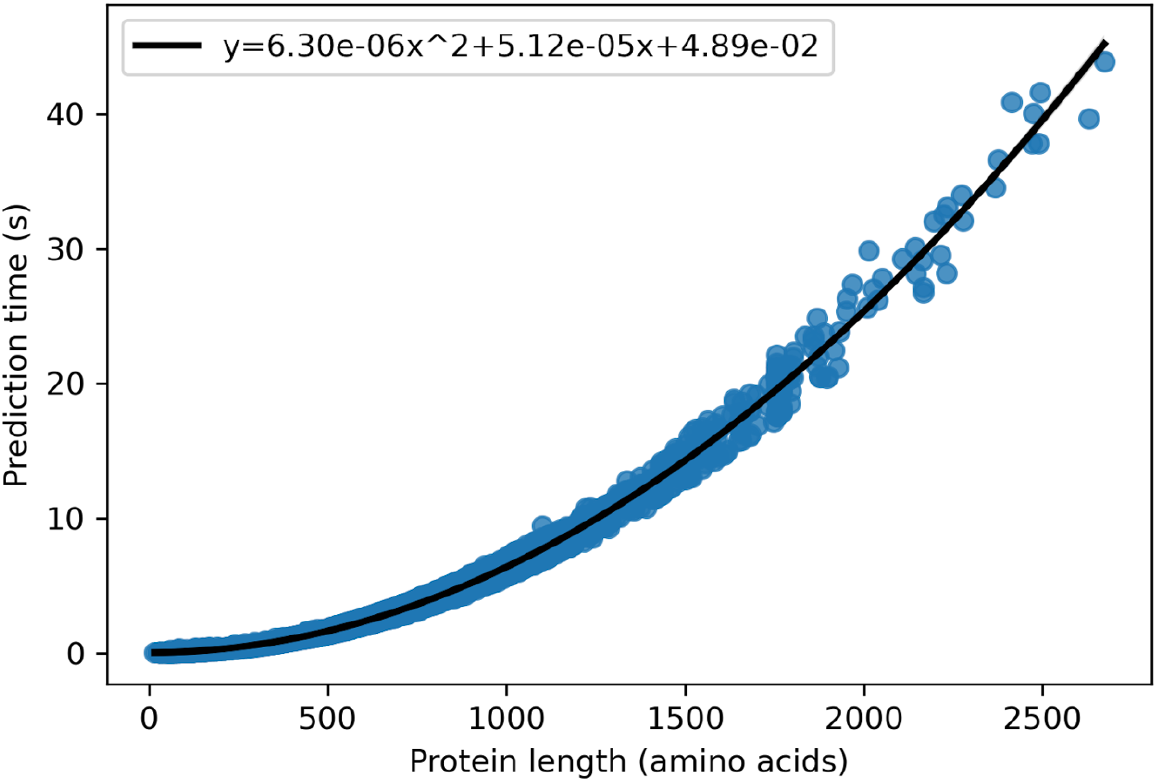
Relationship between protein length and prediction time for SOLeNNoID method on the YEAST v1 AlphaFoldDB dataset. Top left - second order polynomial equation fit to the data using numpy.polyfit.

**Table S2.** Comparison of method performance on the solenoid test set as a binary (solenoid/non-solenoid) classification problem. The metrics used are precision, recall and F1-score for single classes and multi-class MCC as a global metric. Values in bold represent the best performance across methods.

| Precision |  |  |  |  |
| --- | --- | --- | --- | --- |
| Class | SOLeNNoid | TAPO | PRIGSA2 | RDB-lite |
| Non-solenoid | **0.786** | 0.780 | 0.509 | 0.494 |
| Solenoid | 0.870 | 0.855 | **0.954** | 0.892 |
| Recall |  |  |  |  |
| Class | SOLeNNoid | TAPO | PRIGSA2 | RDB-lite |
| Non-solenoid | 0.760 | 0.729 | **0.958** | 0.892 |
| Solenoid | **0.886** | **0.886** | 0.488 | 0.494 |
| F1-score |  |  |  |  |
| Class | SOLeNNoid | TAPO | PRIGSA2 | RDB-lite |
| Non-solenoid | **0.773** | 0.754 | 0.665 | 0.636 |
| Solenoid | **0.877** | 0.870 | 0.646 | 0.636 |
| MCC |  |  |  |  |
|  | SOLeNNoid | TAPO | PRIGSA2 | RDB-lite |
|  | **0.651** | 0.625 | 0.455 | 0.387 |

**Fig. S3.**
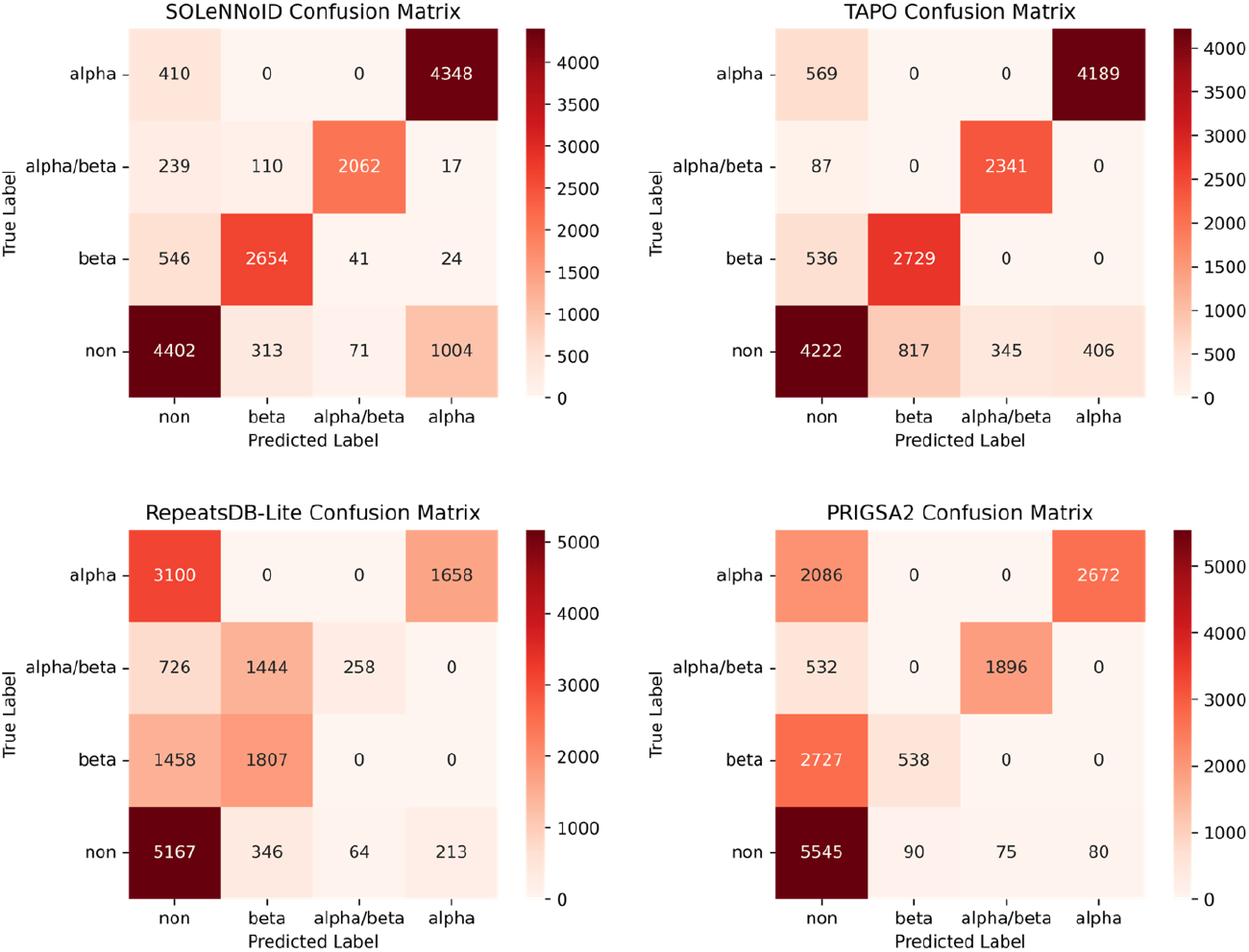
Multi-class confusion matrices for SOLeNNoID, TAPO, PRIGSA2 and RepeatsDB-Lite on the solenoid test set. True labels are shown on the y-axis and predicted labels are shown on the x-axis. The numbers represent the number of protein residues placed within each category of the confusion matrix. A darker red colour indicates a larger number of residues.

**Table S3.** Performance of SOLeNNoID on the whole test set as a multi-class classification problem.

| Class | Precision | Recall | F1-score |
| --- | --- | --- | --- |
| Non-solenoid | 0.935 | 0.912 | 0.924 |
| β-solenoid | 0.860 | 0.813 | 0.836 |
| $\alpha/\beta$-solenoid | 0.948 | 0.849 | 0.896 |
| α-solenoid | 0.770 | 0.914 | 0.835 |
| MCC |  |  |  |
| 0.811 |  |  |  |

**Fig. S4.**
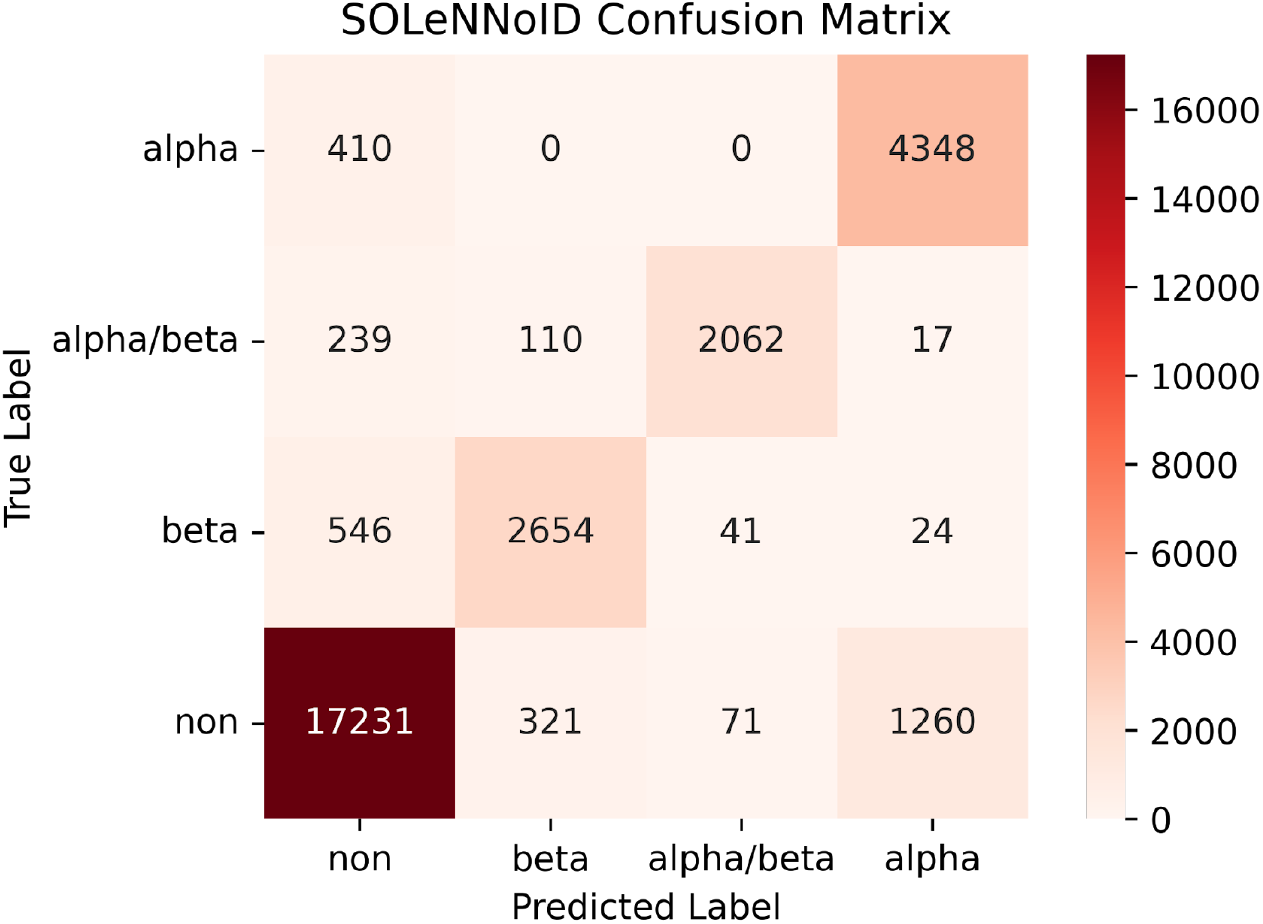
Multi-class confusion matrix for the SOLeNNoID model on the full test dataset. True labels are shown on the y-axis and predicted labels are shown on the x-axis. The numbers represent the number of protein residues placed within each category of the confusion matrix. A darker red colour indicates a larger number of residues.

**Table S4.**
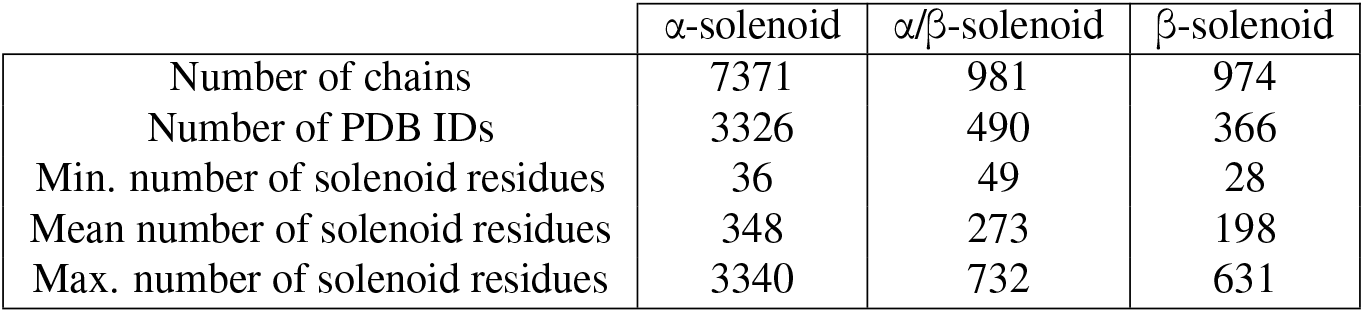
Statistics for PDB entries detected as solenoids by SOLeNNoID.

| | α-solenoid | $\alpha/\beta$-solenoid | β-solenoid |
| --- | --- | --- | --- |
| Number of chains | 7371 | 981 | 974 |
| Number of PDB IDs | 3326 | 490 | 366 |
| Min. number of solenoid residues | 36 | 49 | 28 |
| Mean number of solenoid residues | 348 | 273 | 198 |
| Max. number of solenoid residues | 3340 | 732 | 631 |

